# «The LAMP-CRISPR-Cas13a technique for detecting CBASS-mechanism of phage resistance in bacteria»

**DOI:** 10.1101/2024.12.19.629325

**Authors:** Concha Ortiz-Cartagena, Patricia Fernández-Grela, Lucia Armán, Clara Ibarguren, Lucía Blasco, Daniel Pablo-Marcos, Inés Bleriot, Laura Fernández-García, Felipe Fernández Cuenca, Antonio Barrio-Pujante, Belén Aracil, María Tomás

## Abstract

Antimicrobial resistance (AMR) is an important threat to public health that has led to the development of innovative alternative treatments for bacterial infections, such as phage therapy. However, one of the greatest disadvantages of phage therapy is the generation of phage-resistant bacterial mutants via bacterial defence mechanisms, which are mainly contained in genomic islands (GIs) and controlled by the quorum sensing (QS) network. In this study, 309 pathogenic islands (PAIs) harbouring a total of 22.1 % of proteins related to anti-phage defence (APD) were detected in the genome of 48 K. pneumoniae strains. Several type I and type II CBASS systems were also detected in the genome of the 48 K. pneumoniae strains, but only 2 type II CBASS systems were located in PAIs. We constructed a knockout K. pneumoniae strain, not expressing the cyclase gene from the type II CBASS system present in PAIs, to study the regulatory role of QS in expressing the gene. As the anti-phage CBASS system is an abortive infection (Abi) system, the role of the type II CBASS system in regulating cell viability was assessed. The knockout strain was confirmed by targeting the LAMP-CRISPR-Cas13a technique specifically to the cyclase gene, and the same protocol was also used to detect the gene of the main cyclase in these type I CBASS systems, i.e. APECO1. The study findings demonstrate the regulatory role of the QS network in anti-phage defence systems.

Finally, this is the first work which development an innovative biotechnological application for the LAMP-CRISPR-Cas13a rapid-technique (<2 hours) in optimizing phage therapy by detecting bacterial resistance mechanisms, by predicting the potential inefficacy of a therapeutic phage and thus improving patient prognosis.

## INTRODUCTION

Antimicrobial resistance (AMR) is one of the current main global threats to public health, and it has been predicted that as many as 10 million people could die annually as a result of AMR by 2050 (1–3). Worryingly, the 2022 Global Antimicrobial Resistance and Use Surveillance System (GLASS) report raised concerns about extremely high resistance rates among prevalent bacterial pathogens (4). In this context, carbapenem-resistant Klebsiella pneumoniae is highlighted as a key pathogen against which common antibiotics no longer work. This critical situation has led to increased administration of last-resort drugs, with resistance to these drugs increasing rapidly (5). K. pneumoniae is included in the bacterial priority pathogens list (BPPL) compiled by the World Health Organization (WHO), implying that measures for prevention and appropriate treatment of infections are crucial (6).

Regarding the scarcity of effective treatments for AMR infections, due to the emergence of resistance to all antibiotics used in clinical settings, therapeutic innovations such as phage therapy are becoming more important (7). Phage therapy, the use of bacteriophages (phages, i.e. viruses that infect bacteria) to treat bacterial infections, has several advantages over other alternative treatments (8, 9, 10). However, contact between bacteria and phage leads to the so-called phage-host arms race, and one of the greatest disadvantages of phage therapy is the frequent and rapid generation of phage-resistant bacterial mutants during treatment (11, 12). Anti-phage defence mechanisms in bacteria are mainly based on (i) adsorption resistance, (ii) blockage of the phage infection once adsorbed, (iii) elimination of the viral genetic material (restriction-modification and CRISPR-Cas systems) and (iv) other mechanisms (toxin-antitoxin systems and abortive infection [Abi] systems such as CBASS) (12, 13).

Anti-phage defence systems are commonly contained in clusters of genes within a bacterial genome, known as genomic islands (GIs). These genes appear to have been acquired through horizontal gene transfer (HGT), which rapidly spreads genes that improve bacterial fitness and adaptation (14–16). The genes harboured in GIs are classified as pathogenic islands (PAIs), metabolic islands (MIs), symbiotic islands (SIs) and resistance islands (RIs) (16, 17). Although phage defence systems are frequently clustered together in bacterial genomes in defence islands (DIs), several studies have investigated PAIs containing these systems, such as the phage-inducible chromosomal islands (PICIs) (13, 18). PICIs are PAIs that provide many defence systems in Gram-positive and Gram-negative bacteria, and they also protect and thus benefit the bacterial host from competing phages and other mobile genetic elements (13, 18).

Among the anti-phage defence systems, the cyclic-oligonucleotide-based anti-phage signalling systems (CBASS) are a family of defence systems composed of an oligonucleotide cyclase, which generates signalling cyclic oligonucleotides in response to phage infection. These cyclic oligonucleotides activate an effector that promotes cell death before phage replication is completed, therefore preventing the spread of phages to nearby cells (19). Diverse CBASS defence systems have been identified and are classified according to the composition of the CBASS operon, the activity of the effector protein and the signalling molecule produced by the oligonucleotide cyclase. The type I and II systems are the most common. The type I CBASS system consists of an oligonucleotide cyclase and an effector protein, the so-called “minimal configuration”, and the effector activity is mainly based on the formation of pores in the membrane. In addition to the cyclase-effector framework, the type II CBASS system has 2 ancillary genes (cap2 and cap3) that are involved in expanding the range of phages against which it acts, and the effector protein is usually a phospholipase (19).

Importantly, many bacteria control the expression of several of these defence mechanisms, such as CBASS systems, through the quorum sensing (QS) network, which is defined as a cellular communication process mediated by signalling molecules called autoinducers (AIs) (20, 21). The QS of K. pneumoniae mainly relies on the use of autoinducer-2 (AI-2). AI-2 consists of a group of 4,5-dihydroxy-2,3-pentanedione (DPD) derivatives that can rapidly convert to one another, are involved in interspecies communication and are capable of detecting other autoinducers in the medium such as AI-1 (exogenous N-Acyl homoserine lactones or AHLs) (21, 22). The role of QS in controlling phage defence was demonstrated in a previous study, which showed that inhibition of QS reduces expression of the CBASS proteins and finally to an improvement of the phage infection (20). Under this premise, the strategy of counteracting CBASS phage defence systems may be a useful way of enhancing the therapeutic outcomes of phages in clinical settings.

Detection of the anti-phage systems harboured in a targeted clinical strain can be used to overcome bacterial resistance to phage, hence enabling the application of personalized phage-therapy treatments that improve patient prognosis. In this context, CRISPR-Cas systems are potentially promising tools, as they have yielded successful results regarding the development of detection protocols with high levels of sensitivity and specificity (23–25). These diagnostic techniques have previously been used to detect several microorganisms and also resistance genes (26, 27).

In this study, we analysed the anti-phage defence genes contained in the PAIs detected in the genome of 48 K. pneumoniae isolates. We also conducted an epidemiological study of the CBASS systems located in these strains, focusing on the QS-based regulation of the type II CBASS anti-phage defence system in a carbapenemase-producing K. pneumoniae clinical strain. Finally, we broadened the range of biotechnological applications of CRISPR-Cas technology, by applying our previously developed diagnostic protocol based on the CRISPR-Cas13a system to detect anti-phage resistance genes in bacteria.

## MATERIAL AND METHODS

### IN SILICO ANALYSIS

Genomes in contigs of 48 K. pneumoniae clinical isolates (Table 1), previously sequenced by Illummina miseq (Bioproject codes listed in Table 1) (28, 29), were studied. Contigs were assembled through an aleatory number of “Ns” to compress the genomes into single sequences, which were then reannotated using the PROKKA software tool (30), in both cases with the Linux OS.

**Table 1.** Bacteria used in the present study. ^a^ BioSample code of complete genomes included in the European BioProject PRJNA565865. ^b^ BioSample code of complete genomes included in the European BioProject PRJEB10018.

| Hospitals | National Centre of Microbiology<br>(Madrid, Spain)<sup>a</sup> | Virgen Macarena University Hospital<br>(Seville, Spain)<sup>b</sup> |
| --- | --- | --- |
| Strain | 1ST405OXA48 | K2535S |
|  | 2ST15VIM1 | K2551S |
|  | 3ST11OXA245 | K2597S |
|  | 4ST437OXA245 | K2691S |
|  | 5ST16OXA48 | K2707S |
|  | 6ST101KPC2 | K2715S |
|  | 7ST147VIM1 | K2783S |
|  | 8ST11VIM1 | K2791S |
|  | 9ST846OXA48 | K2982S |
|  | 10ST340-VIM1 | K2983S |
|  | 12ST13OXA48 | K2984S |
|  | 13ST512KPC3 | K2986S |
|  | 14ST15OXA48 | K2989S |
|  | 15ST11OXA48 | K2990S |
|  | 16ST258KPC3 | K3318S |
|  | 17ST899OXA48 | K3320S |
|  | 18ST974OXA48 | K3321S |
|  | SCISP2CS | K3322S |
|  | SCISP4CS | K3323S |
|  |  | K3324S |
|  |  | K3325S |
|  |  | K3416S |
|  |  | K3509S |
|  |  | K3571S |
|  |  | K3572S |
|  |  | K3573S |
|  |  | K3574S |
|  |  | K3575S |
|  |  | K3579S |
|  |  | K3667S |

### IDENTIFICATION AND ANALYSIS OF GENOMIC ISLANDS

Genomes from 48 K. pneumoniae clinical isolates were analysed using GIPSy software (31) to predict the genomic islands (GIs) they harboured. The GIs were classified into pathogenicity islands (PAIs), metabolism islands (MIs), symbiotic islands (SIs) and resistance islands (RIs) by using GIPSY software, with the genomes in GBK format files as input and the genome of K. pneumoniae strain MGH78578 (Genbank: NC_009648) as the reference genome. The sequences of the predicted PAIs were extracted from their respective genomes; the “Ns” were removed and the sequences were annotated using the PROKKA software tool, in all cases with the Linux OS. The PAIs sequences were analysed, using the HHMER (32) and HHPred (33) servers, along with the NCBI conserved domains database (34), to identify and thus classify the proteins contained in them, as follows: (i) metabolic proteins; (ii) defence, resistance and virulence proteins (DRV); (iii) DNA metabolism proteins; (iv) mobile genetic elements (MGE) and (v) unknown proteins. Anti-phage defence (APD) genes were also detected using PADLOC, a web server for the identification of antiviral defence systems in microbial genomes, and the defence sequences were analysed using HHMER and HHPred servers (35).

### CYCLASE GENE KNOCKOUT STRAIN

#### Strains, plasmids and reagents

The K. pneumoniae strain 10ST340-VIM1 was selected from the bacterial strain collection (Table 1), to construct a knockout of the cyclase gene (10ST340-VIM1Δcyclase) present in a type II CBASS system, by using the E. coli λ-Red system, with some modifications (36). The homologous regions, required for the recombination step in the target genome, were of length 300 bp, and therefore the length of the knockout cassette ends was increased to 300 bp. Plasmids and primers used for this goal are listed in Table 2.

**Table 2.**
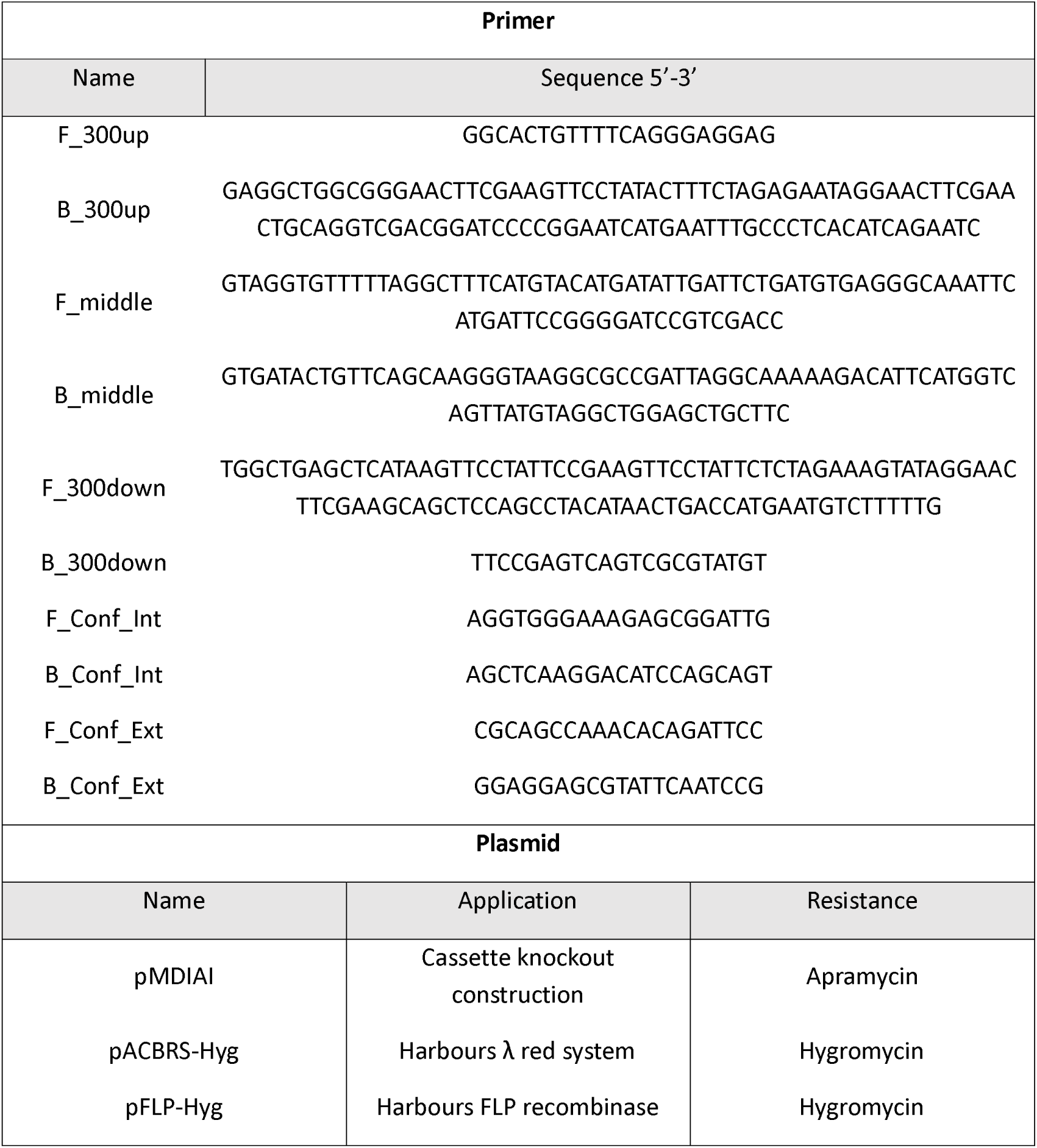
Oligonucleotides and plasmids used to construct the knockout strain in the present study.

Bacterial cultures were grown in Luria Bertani (LB) broth (10 g/L tryptone, 5 g/L yeast extract, and 10 g/L NaCl) or on LB agar (LB broth supplemented with 20 g/L of agar). Hygromycin and apramycin were used to select transformant strains at concentrations of 100 µg/mL and 50 µg/mL, respectively.

The cyclase gene was deleted from the K. pneumoniae 10ST340-VIM1 strain by the above-mentioned method (36). Briefly, the cassette knockout containing the apramycin resistance gene was electroporated into the K. pneumoniae 10ST340-VIM1 strain already transformed with plasmid pACBSR-Hyg, which expresses the λ-Red system and a hygromycin selection marker. Bacteria were incubated overnight on LB Agar supplemented with apramycin, and correct insertion of the cassette knockout was confirmed by PCR (F_300up and B_Conf_Int). The K. pneumoniae 10ST340-VIM1 derivative strain was grown on LB agar with apramycin (50 μg/mL), at 37 °C for 3 days, before being incubated on LB agar with apramycin (150 μg/mL) and LB agar with hygromycin (100 μg/mL), overnight at 37 °C, to test for loss of the helper plasmid pACBSR-Hyg. For removal of resistance markers, cells were electroporated with pFLP-Hyg and grown overnight at 30 °C. The mutants were then incubated at 43 °C to remove the apramycin resistance from the inserted knockout cassette. Apramycin-susceptible mutants were selected to confirm deletion of the cyclase gene by PCRs (F_300up/B_300down and F_Conf_Int/B_Conf_Int) and Sanger sequencing (Macrogen®).

### INDUCTION OF EXPRESSION OF THE CBASS BY QS

Expression of effector I was quantified in the presence of 20 µM of AHL C6-HSL in both the 10ST340-VIM1 and 10ST340-VIM1Δcyclase K. pneumoniae strains. For this purpose, 40 μL of an overnight culture of the corresponding K. pneumoniae strain was inoculated in 4 mL flasks of LB-broth containing 20 µM C6-HSL (two biological samples of each strain). The flasks were then incubated at 37 °C, with shaking at 180 rpm, until the OD_600_ (optical density measured at wavelength of 600 nm) reached 0.3.

Duplicate aliquots of 1 mL were removed from each flask for subsequent RNA extraction with the High Pure RNA Isolation Kit (Roche). Extraction was conducted following the manufacturer’s instructions. The RNA extracted from each sample was measured with a Nanodrop spectrophotometer (NanoDrop Technologies) and adjusted to 50 ng/μL with nuclease-free water for use in qRT-PCR (LightCycler® 480). Specific primers and the corresponding UPL probe for both the effector I and recA (Gomes et al., 2018) genes (Table 4), were used in this assay, conducted with the LightCycler® 480 Control Kit (Roche). A Student’s t-test (GraphPad Prism 9.0.0) was used to determine any statistically significant differences (P-value < 0.05) in gene expression.

### CELL VIABILITY IN THE PRESENCE OF OVEREXPRESSION OF CBASS

Cell proliferation and cell viability levels in both 10ST340-VIM1 and 10ST340-VIM1Δcyclase K. pneumoniae strains cultured in the presence of 20 µM of C6-HSL (to overexpress the CBASS) were determined by an assay based on the WST-1 reagent, a red tetrazolium salt that turns yellow when cleaved to formazan by mitochondrial dehydrogenases (WST-1 Assay Protocol for Cell Viability, MERCK®). Briefly, overnight cultures of the corresponding K. pneumoniae strains were diluted 1:100 in LB-broth (two biological samples of each strain) and the flasks were incubated at 37 °C, with shaking at 180 rpm, until the OD_600_ reached 0.2. The cultures were then diluted to OD_600_ 0.02 and added to the wells of 96-well plates and made up to a final volume of 100 µL supplemented with 20 µM of C6-HSL. The plates were incubated at 37 °C, with shaking at 180 rpm for 24 h, and the OD_600_ was again read in a microplate reader (NanoQuant). Dilutions were made in new 96-well plates to adjust the OD_600_ to 0.02, and 10 µL of WST-1 reagent was then added to the plates, which were incubated at 37 °C in absence of light for 1.5 h. To quantify proliferation and viability, the OD was measured at a wavelength of 480 nm (OD_480_). A Student’s t-test (GraphPad Prism 9.0.0) was used to determine any statistically significant differences (P-value < 0.05) in proliferation levels. In this assay, negative controls consisted of LB without strains nor C6-HSL supplementation.

### DETECTION OF CBASS BY CRISPR-CAS13

The LAMP-CRISPR-Cas13a detection protocol, previously described by our research group (26, 27), was used to detect the strains harbouring type I and type II CBASS systems from among the 48 the K. pneumoniae isolates. In order to identify type I CBASS systems, the LAMP-CRISPR-Cas13a technique was targeted to the cyclase APECO1 gene, located in several type I CBASS systems; the presence of type II CBASS systems was determined by identifying the cyclase gene contained in both type II CBASS systems detected in the PAIs. For this purpose, genomic DNA was extracted from the respective strains by the proteinase K-heat inactivation protocol (PK-HID) (37). Briefly, a LAMP DNA amplification reaction (WarmStart® LAMP Kit (DNA & RNA), NEB, Ipswich, MA, USA) was performed for cyclase gene amplification, containing 5 µl of extraction, 12.5 µl of WarmStart LAMP 2x Master Mix and 2.5 µl of Primer Mix 10x (FIP/BIP 16 µM, F3/B3 2 µM, LOOPF/LOOPB 4 µM, stock 10X) adjusted to a final volume of 25 µl with RNase-free water and incubated at 65 °C for 1 hour. Following, a Leptotrichia wadei, LwaCas13a, endonuclease (GenCRISPR™ Cas13a (C2c2) Nuclease, GenScript) detection was assessed, using 2 µl of cleavage buffer 10X (GenCRISPR™ Cas13a (C2c2) Nuclease, GenScript), 0.5 µl of dNTPs (HiScribe™ T7 Quick High Yield RNA Synthesis Kit, NEB), 0.5 µl of T7 polymerase (HiScribe™ T7 Quick High Yield RNA Synthesis Kit, NEB), 20 U of RNase murine inhibitor (NEB), 25 nM of LwaCas13a endonuclease, 50 nM of crRNA (IDT), 1000 nM of reporter (GenScript) and 5 µl of LAMP amplicon, adjusted to a final volume of 20 µl with RNase-free water and incubated at 37 °C for 30 min (Table 3). Finally, the test results were revealed using HybriDetect lateral flow test strips (Milenia Biotec, Giessen, Germany), mixing 20 µl of collateral cleavage activity-based detection product with 80 µl of assay buffer supplemented with 5 % polyethylenglycol in a 96-well plate where strip tests were placed. The test results are visible to the naked eye after 2-3 minutes. In summarize the duration of the development of the LAMP-CRISPR-Cas13a rapid-technology is less than 2 hours.

**Table 3.**
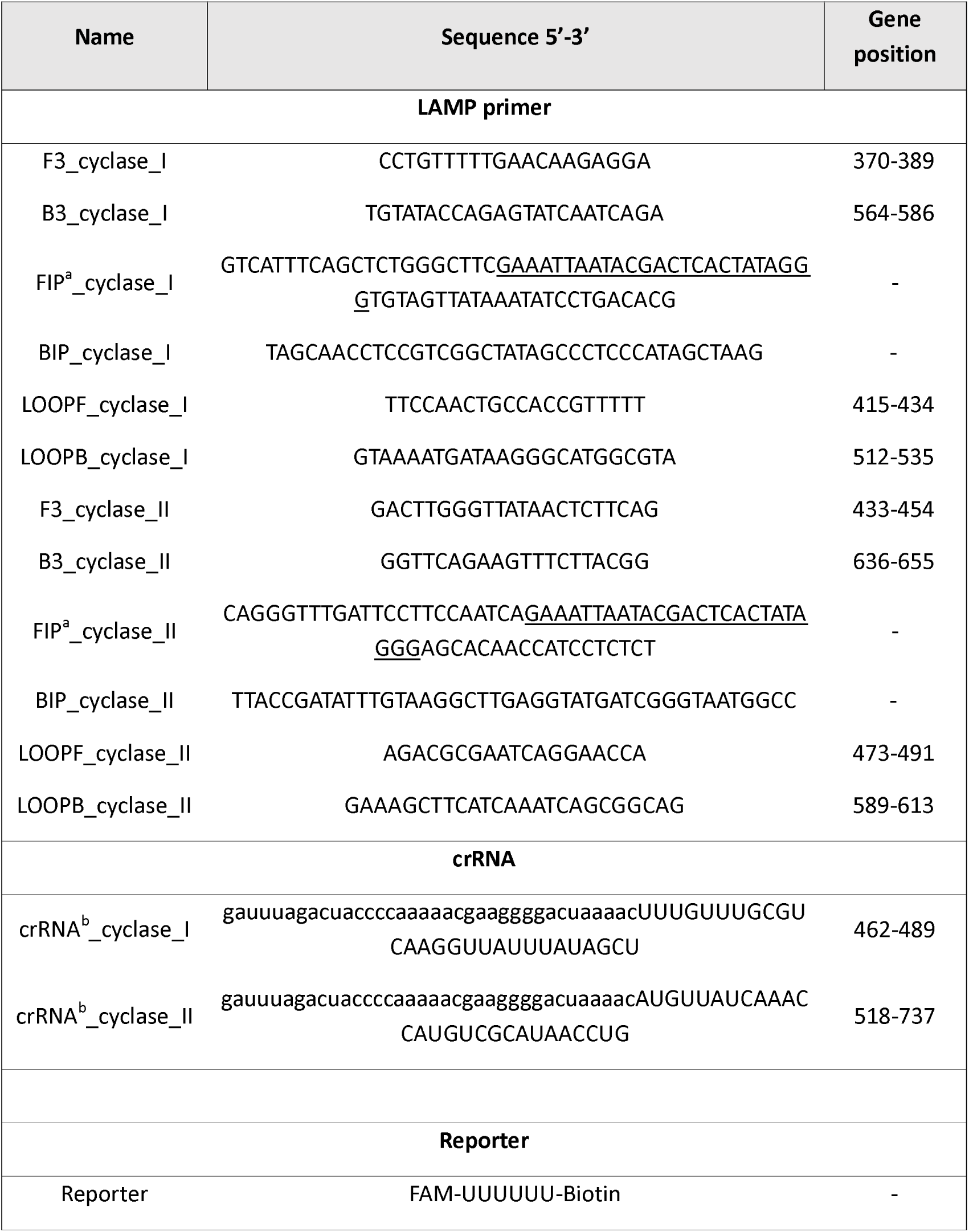
Sequences of primers, crRNA and the reporters used for LAMP-CRISPR-Cas13a detection. All primers were supplied by IDT, and reporter was supplied by GenScript. ^a^Underlined letters indicate the T7 promoter sequence. ^b^Lower-case letters indicate the scaffold sequence.

**Table 4.**
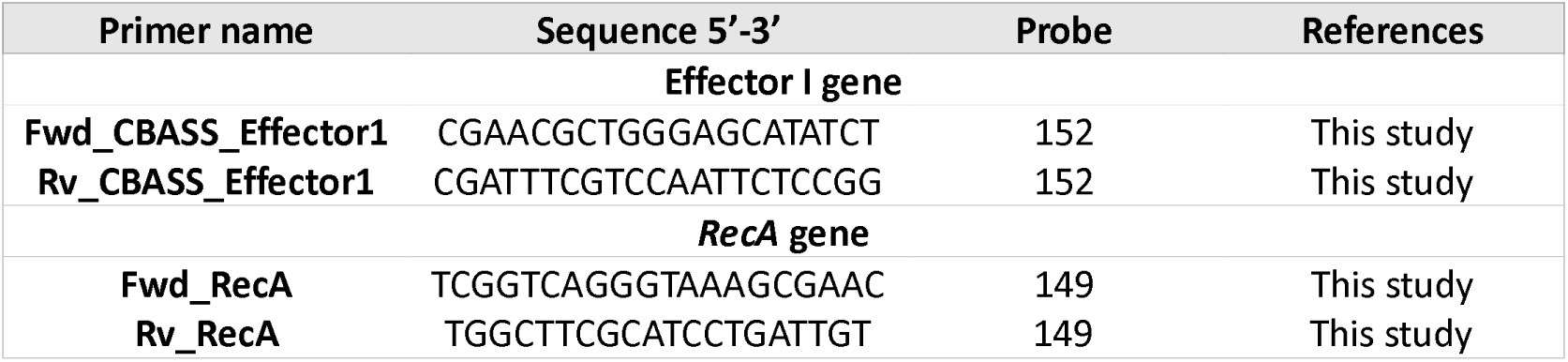
Sequences of oligonucleotides and probes used for quantitative expression of the effector I gene.

## RESULTS

### PAIS AND SEARCH ANALYSIS

Analysis of the genomes of 48 clinical isolates of K. pneumoniae led to the identification of a total of 309 regions with a high probability of being PAIs. Among these, 17 PAIs were detected in more than one K. pneumoniae strain, and the sequences were analysed bioinformatically to determine the function of the proteins harboured in them, as greater spread could be related to important roles in bacterial fitness and adaptation (Figure 1A). The results showed that 31 % of the proteins were related to mobile genetic elements (MGE), 26 % to defence-resistance-virulence (DRV), 13 % to DNA metabolism, 11 % to metabolism and 19 % to unknown function. Furthermore, 24.4 % of these proteins were involved in APD, and 46 % of these corresponded to the DRV functional group, but also 25 % to MGE, 12 % to DNA metabolism, 9 % to metabolism and 8 % were of unknown function. Notably, 90.5 % of the APD proteins in MGE were proteins of phage origin. Finally, organization of the APD proteins in the different APD systems revealed that most of these proteins are involved in surface modification (28.6 %), toxin-antitoxin (TA) (22.9 %) and restriction-modification (RM) (21.4 %) systems. We also identified several complete defence systems: 12 surface modification, 2 TA, 2 RM, 2 CBASS, 1 Gabija, 1 Dynamin, 1 Thoeris and 3 putative systems (Figure 1B).

**Fig 1.**
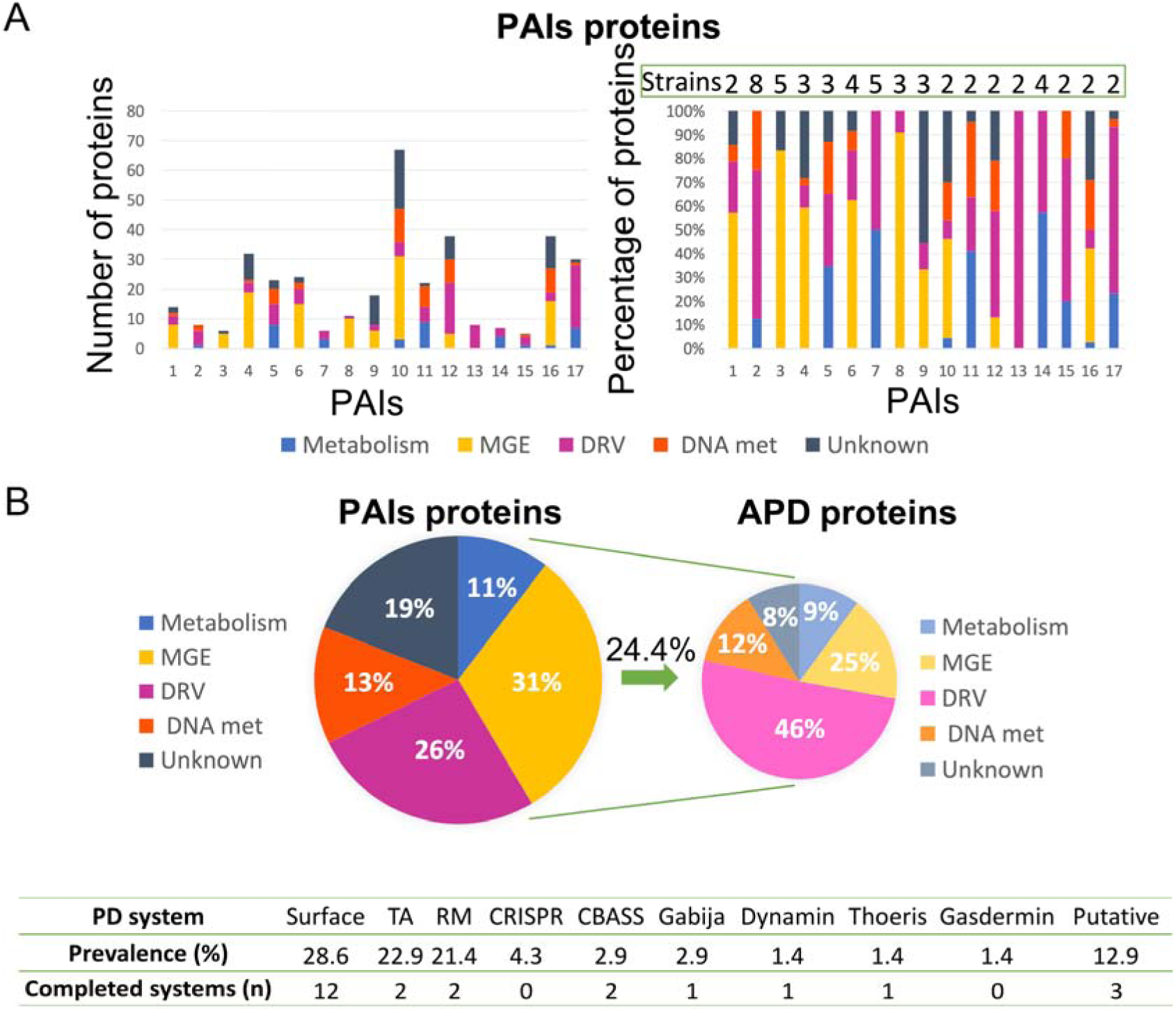
Results of the bioinformatic analysis of proteins contained in 17 Klebsiella pneumoniae PAIs. A: Classification of proteins in functional groups, showing number of proteins (left) and percentages of proteins (right) in each PAI; the number of K. pneumoniae strains containing each PAI is also shown (right). B: Global percentages of groups of functional proteins in PAIs (left); ADP proteins classified in the different functional groups (right); and table containing the PD systems identified in these PAIs, their prevalence and the number of the completed systems found (below).

The complete genome of the K. pneumoniae strains with CBASS systems harboured in PAIs, 10ST340-VIM1 and K3325S were exhaustively analysed by PADLOC, along with the genomes from the other 46 K. pneumoniae strains, to identify CBASS systems not contained in PAIs. The genomic structure of the CBASS systems in both 10ST340-VIM1 and K3325S strains was similar (Table 5) and only these two CBASS systems (11 %) were located in PAIs (Table 6). Regarding classification of the CBASS systems detected, 63 % belonged to type I and 37 % to type II. None of the type I CBASS systems were contained in PAIs, and two of the three type II CBASS systems were located in PAIs of the 10ST340-VIM1 and K3325S strains. Analysis of the components of these CBASS systems revealed that the cyclase APECO1 gene was present in 13 of the 15 (87 %) type I CBASS systems detected but not in any of the type II CBASS systems. Regarding the effector protein, the pore-forming function effector appeared in all type I and in one of three type II (33 %) CBASS systems; however, phospholipase and nuclease effectors were only located in type II CBASS systems (Table 6). Finally, the cyclic-nucleotide signal was GMPAMPc in all of the CBASS systems analysed (Table 6).

**Table 5.**
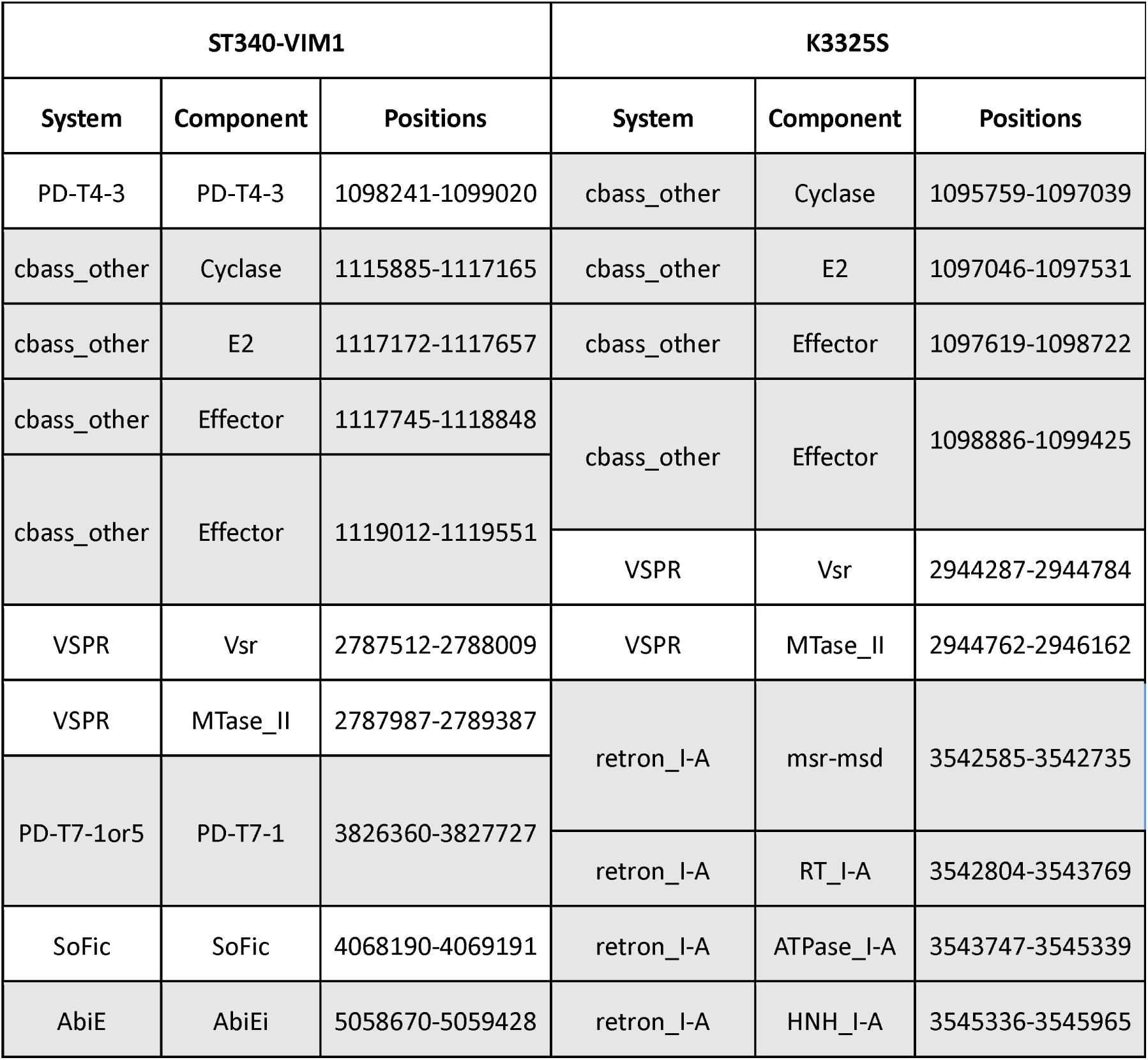
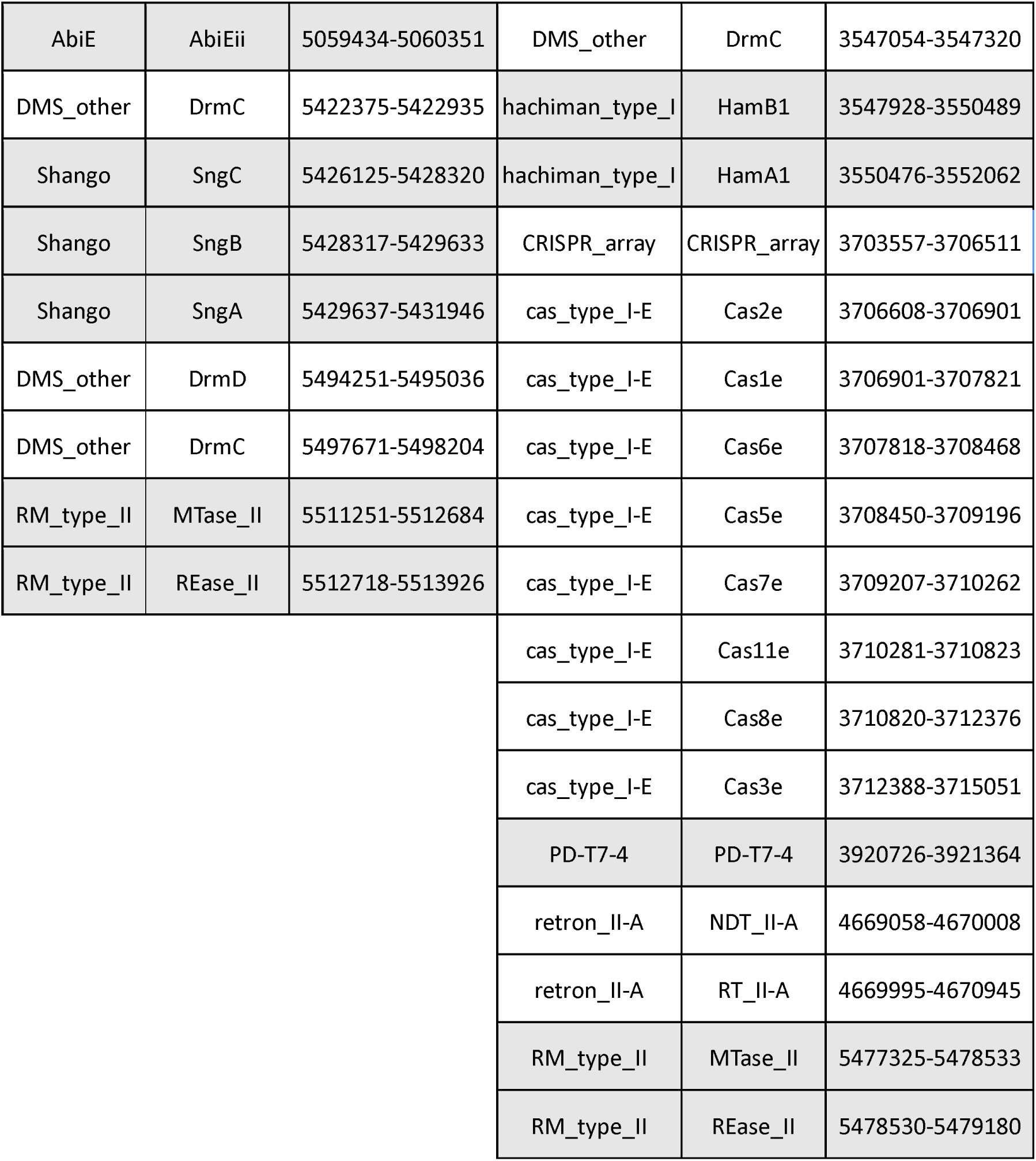
Results of the bioinformatic analysis of the APD systems harboured in 10ST340-VIM1 and K3325S K. pneumoniae strains.

**Table 6.**
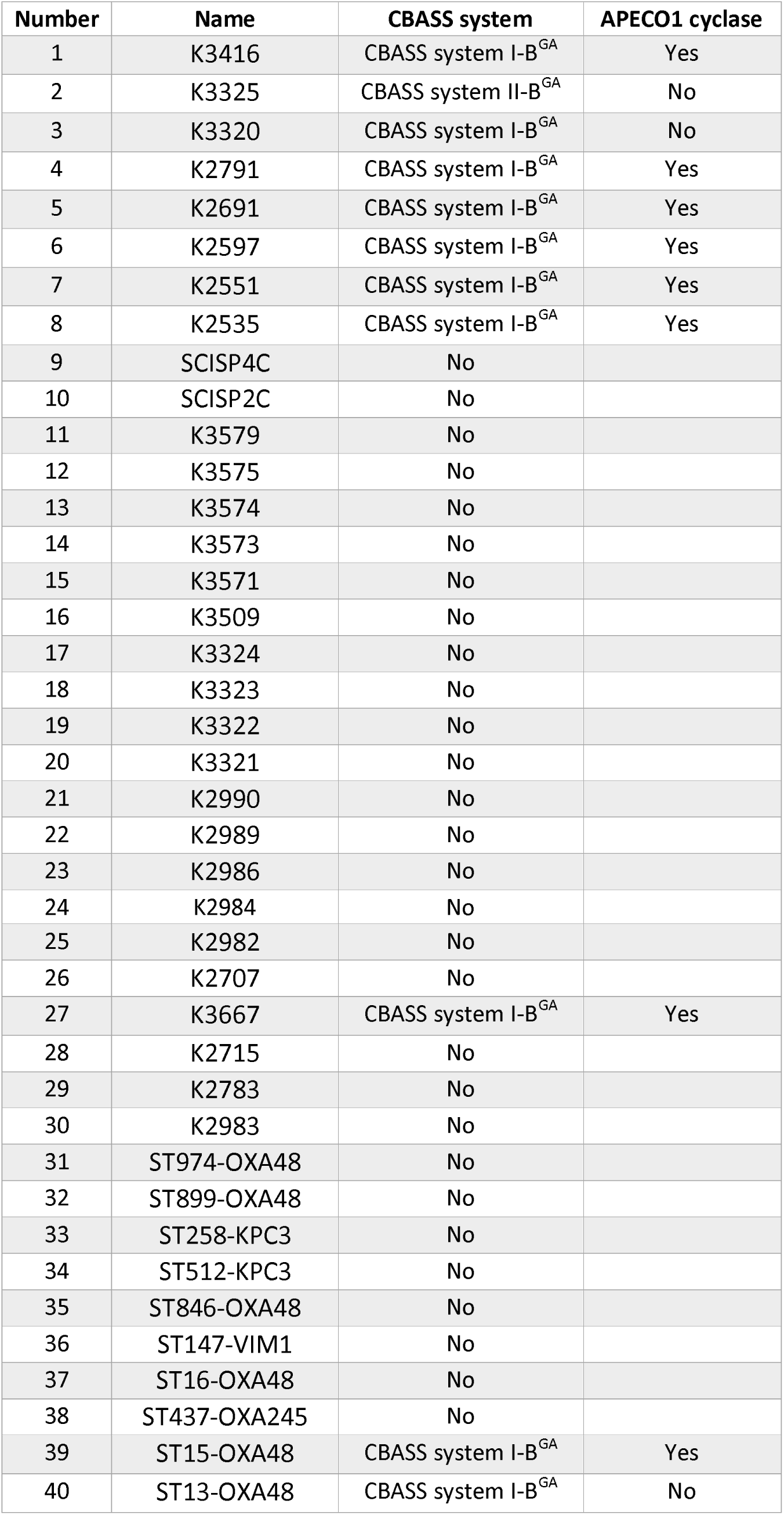
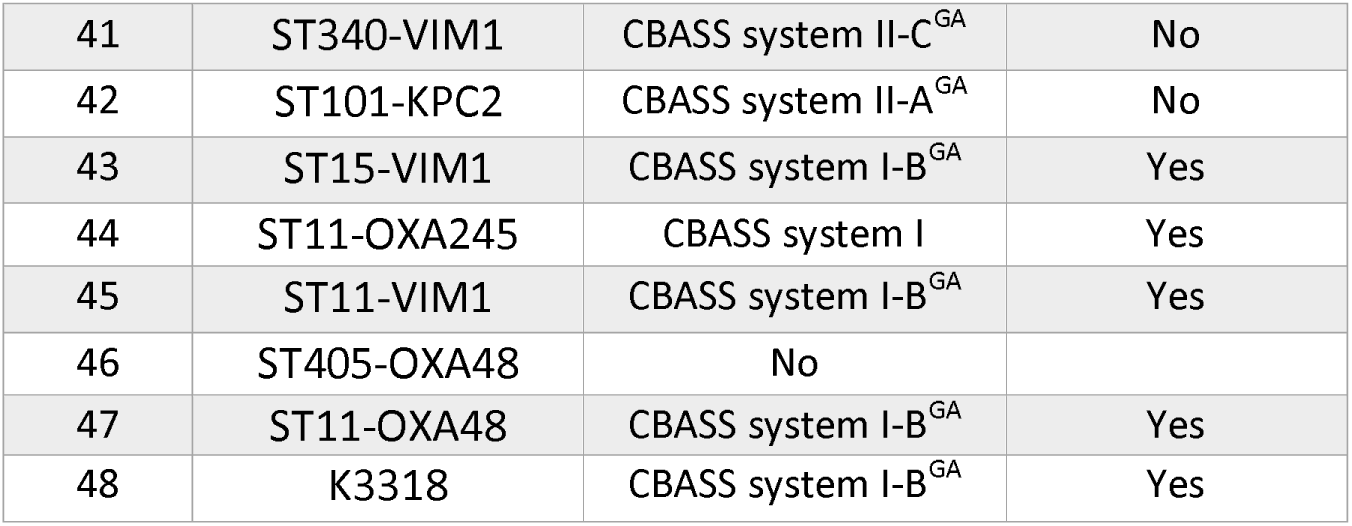
Results of the bioinformatic analysis of the CBASS systems contained in the 48 Klebsiella pneumoniae strains. A: Type I CBASS systems with a phospholipase as effector; B: Type I CBASS systems with a pore-forming effector activity; C: Type I CBASS systems with an endonuclease effector. ^GA^ Cyclic GMP-AMP second messenger signal.

### REGULATORY ROLE OF QUORUM SENSING IN CBASS SYSTEM EXPRESSION

In order to study the regulatory role of quorum sensing in expression of the type II CBASS system, we constructed a 10ST340-VIM1Δcyclase K. pneumoniae strain, and deletion of the gene was consecutively confirmed by the LAMP-CRISPR-Cas13a technology (Figure 2). Quantitative study of the type II CBASS system expression under QS activation conditions was conducted by measuring effector I gene expression levels in the presence of the AHL C6-HSL, in both 10ST340-VIM1 and 10ST340-VIM1Δcyclase K. pneumoniae strains. The results revealed overexpression of the anti-phage defence system in the 10ST340-VIM1 K. pneumoniae strain (1.23) (Table 7). By contrast, the level of expression of the type II CBASS system in the 10ST340-VIM1Δcyclase K. pneumoniae strain (0.92) was similar to that of the recA housekeeping gene (Table 7).

**Fig 2.**
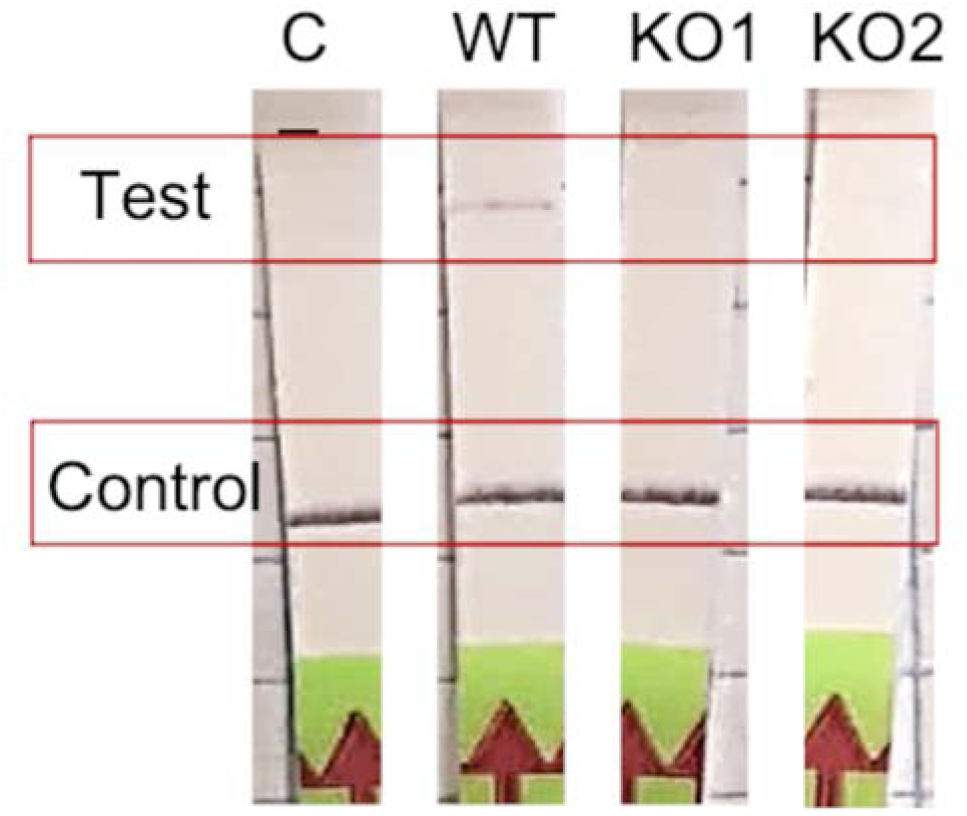
Results of the cyclase gene knockout construction by λ-Red protocol. Confirmation of the deletion of the cyclase gene in two mutants by LAMP-CRISPR-Cas13a protocol.

**Table 7.** Results of the quantitative expression of the effector I gene.

|  | Condition | Cp CBASS |
| --- | --- | --- |
| **WT** | Hex 20 | 1.23 |
| **KO** | Hex 20 | 0.92 |

### INVOLVEMENT OF THE PAI CBASS DEFENCE SYSTEM IN BACTERIAL VIABILITY LEVELS

The role of CBASS (an Abi defence system) in bacterial viability was studied using an assay based on the WST-1 reagent. With this aim, cellular respiration levels in the 10ST340-VIM1 and 10ST340-VIM1Δcyclase K. pneumoniae strains were compared, using the AHL C6-HSL to induce the CBASS system. There was no significant difference in the viability of the 10ST340-VIM1Δcyclase K. pneumoniae strain with and without the autoinducer C6-HSL (Figure 3B), in contrast to the 10ST340-VIM1 K. pneumoniae strain, which showed a significant reduction in cell respiration when treated with AHL C6-HSL (Figure 3A). By contrast, the viability of the 10ST340-VIM1 K. pneumoniae strain was significantly higher than that of the 10ST340-VIM1Δcyclase K. pneumoniae strain (Figure 3C). Finally, the viability of both strains was very similar in the presence of AHL C6-HSL, except in one of the 10ST340-VIM1 K. pneumoniae strains that grew more than the other 10ST340-VIM1 and 10ST340-VIM1Δcyclase K. pneumoniae strains (Figure 3D).

**Fig 3.**
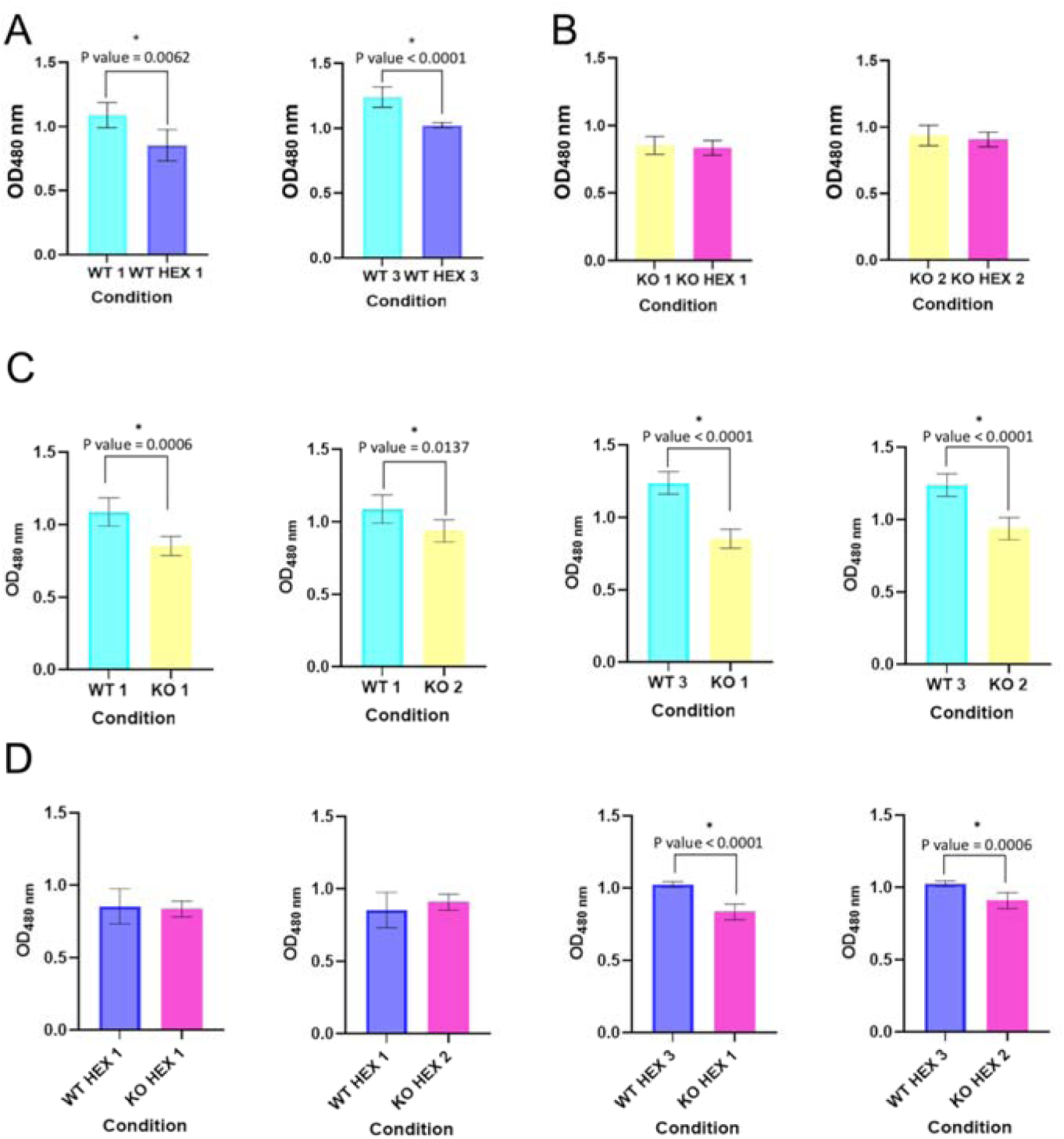
Cell viability in the presence and absence of AHL C6-HSL in both 10ST340-VIM1 and 10ST340-VIM1Δcyclase K. pneumoniae strains. A. Comparison of cell viability in 2 10ST340-VIM1 biological replicates. B. Comparison of cell viability in 2 10ST340-VIM1Δcyclase biological replicates. C. Comparison of cell viability in the 10ST340-VIM1 and 10ST340-VIM1Δcyclase strains in the absence of AHL C6-HSL. D. Comparison of cell viability in the 10ST340-VIM1 and 10ST340-VIM1Δcyclase strains in the presence of AHL C6-HSL.

### DETECTION BY LAMP-CRISPR-CAS13A

The LAMP-CRISPR-Cas13a technique produced very good results regarding detection of the cyclase APECO1 present in several type I CBASS systems (n=13) and the cyclase contained in the type II CBASS systems located in PAIs (n=2). The technique was able to detect the cyclase APECO1 gene in all of the K. pneumoniae strains containing it, except strain number 1. It also produced negative results for the K. pneumoniae strains without CBASS systems and with type II CBASS systems, and for the strains containing type I CBASS systems with a different cyclase (Figure 4A). Importantly, application of a bioinformatic assay to the type I CBASS system in strain 1 indicated that the cyclase APECO1 located in this system had only 7 nucleotides in common with the crRNA molecule, required for Cas13a detection. In addition, the technique successfully detected the presence of the cyclase in the type II CBASS systems contained in PAIs in the 10ST340-VIM1 and K3325S K. pneumoniae strains (Figure 4B). Based on the results obtained, we estimated that the technique exhibits 92.3 % sensitivity and 100 % specificity (99 % confidence interval [CI]) for detection of the cyclase APECO1 gene, as well as 100 % sensitivity and specificity (99 % confidence interval [CI]) for identification of the cyclase gene contained in the type II CBASS systems located in PAIs (Figure 4C).

**Fig 4.**
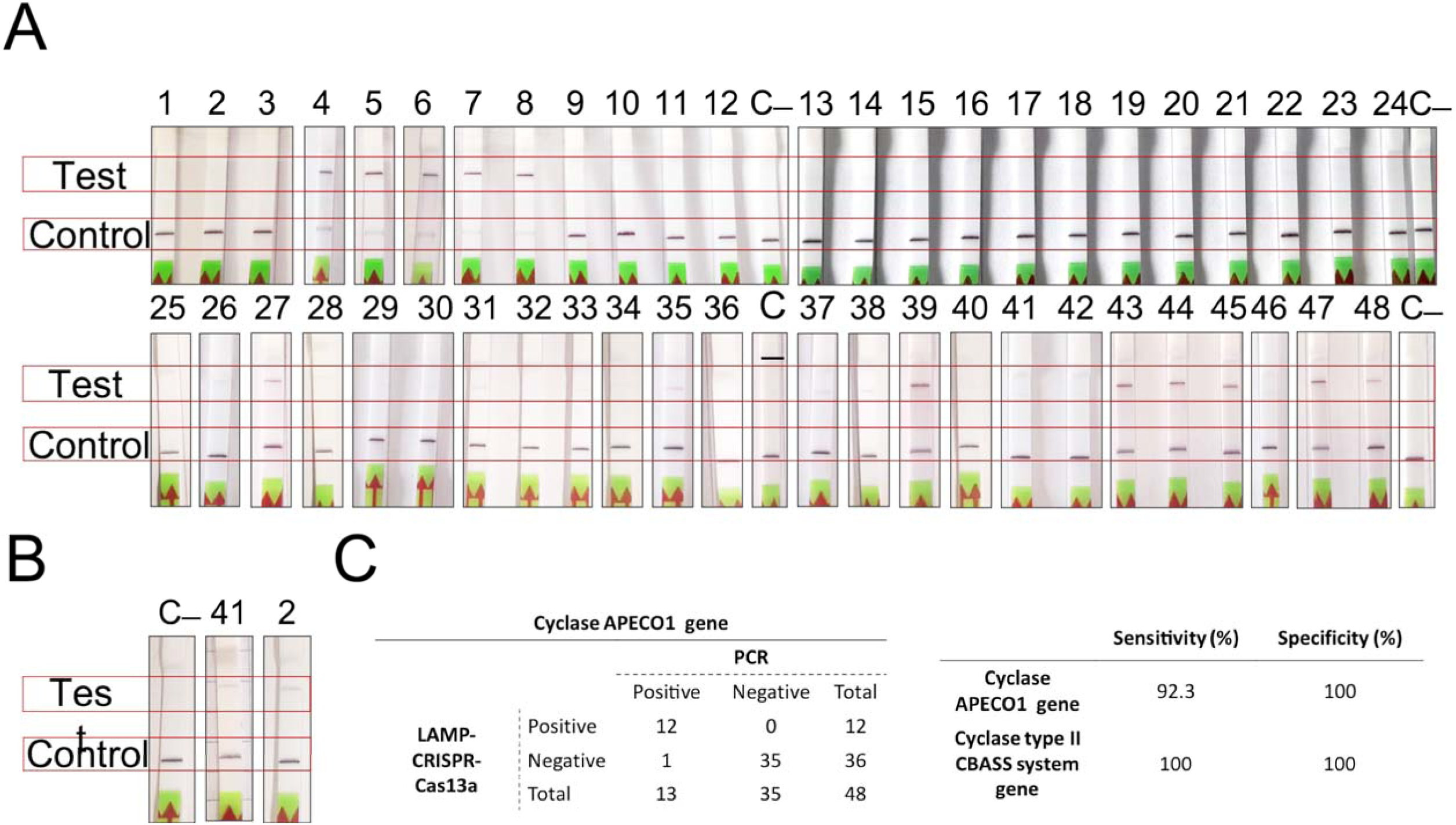
Results of the detection of the type I and II CBASS systems by the LAMP-CRISPR-Cas13a technology. A. Strips for detection of the K. pneumoniae strains harbouring the APECO1 gene; B. Strips for detection of the K. pneumoniae strains containing the cyclase gene in the type II CBASS system; C. APECO1 gene detection (Figure 4A) and table showing the specificity and sensitivity of the LAMP-CRISPR-Cas13a technique, determined by processing data in A and B.

## DISCUSSION

The results of the in silico analysis support the possibility that GIs are essential elements in the HGT for the optimization of bacterial fitness and adaptation, as more than 50 % of the genes found in the 309 PAIs were mobile genetic elements (MGE) or were involved in defence, resistance or virulence (DRV) (Figure 1A). More than 20 % of the proteins contained in the 309 PAIs were related to a single defence mechanism, i.e. anti-phage defence (Figure 1B) (13). This finding is consistent with those of Fillol-Salom et al. (2022), who demonstrated that PICIs (a type of PAIs) are also responsible for transfer of defence genes.

Study of the complete anti-phage defence systems harboured in the 17 most widespread PAIs revealed that these PAIs may be more important for bacterial fitness than previously considered, as the most prevalent anti-phage systems detected have a low cost to bacterial fitness (Figure 1B). Indeed, we did not detect any complete CRISPR-Cas system, in accordance with the increasingly prevalent idea that this anti-phage defence system has a high cost for bacterial fitness, thus explaining its unexpectedly low prevalence across the bacterial domain (38). The cell-surface modification and the defence systems harboured in prophages were notable findings, as preventing phage adsorption is the first line of defence in bacteria, and the main representative system. Prophages are widely spread and provide bacterial defence systems, which explains why almost all MGEs harbouring APD proteins are prophages (39, 40). We also detected other common anti-phage systems: toxin-antitoxin, restriction-modification and CBASS anti-phage defence systems. Abi systems have been detected in a wide variety of microorganisms, although their abundance is difficult to assess owing to the high level of diversity (41).

Analysis of the CBASS systems contained in the 48 genomes of the K. penumoniae strains identified almost twice as many type I CBASS systems as type II CBASS systems. Importantly, the fact that only type II CBASS systems were found in PAIs sequences may indicate that this type of CBASS system is more important for bacterial fitness. This may also explain why the cyclase APECO1 is present in a high proportion of all type I CBASS systems (87 %), while the presence of the cyclase involved in type II systems is more variable. According to the aforementioned explanation, the pore-forming effector function was homogeneously observed in the type I CBASS systems, but the effector harboured in the type II CBASS systems may be related to pore-forming or to phospholipase enzymatic function. Finally, the second signal GMPAMPc seems to be widespread, as it is contained in all the CBASS systems detected.

Since being described in 2020 by Cohen et al., CBASS systems have been detected in the genomes of many prokaryotes, which demonstrates their widespread importance in enabling bacteria to resist phage replication (42). The functional study of the type II CBASS system located in a PAI harboured in the 10ST340-VIM1 K. pneumoniae strain was assessed, due to the role of the PAIs in spreading resistance mechanisms and the fact that the 10ST340-VIM1 isolate contained fewer APD systems than the K3325S strain. The study focused on regulation by the QS network, as well as on the role of the CBASS systems in the proliferation and cellular viability levels. The qRT-PCR assay showed that induction of the QS by AHL C6-HSL led to overexpression of the CBASS system in the 10ST340-VIM1 K. pneumoniae strain, indicating that detection of a threat to the bacterial community via the QS network provokes a signal that finally leads to activation of defence systems, some involved in anti-phage resistance, as previously described by Barrio-Pujante et al. (20) (Figure 2). We also demonstrated that stimulation of the QS network causes a significant reduction in bacterial growth rates when the strain harbours a functional CBASS system, corroborating the aforementioned findings (Figure 3A), as CBASS is an Abi defence system. However, the lower growth rate of the 10ST340-VIM1Δcyclase K. pneumoniae strains suggests that the APD system may be involved in other mechanisms that improve bacterial fitness when growth conditions are suitable (Figure 3C).

Given the importance of new therapeutic strategies like phage therapy, an accurate protocol for rapid detection of mechanisms of bacterial resistance to phages, such as CBASS systems, may be essential for the development of effective phage cocktails or combinations with other therapeutic alternatives, to improve patient prognosis. We have demonstrated the importance of the LAMP-CRISPR-Cas13a technique for detecting the most widespread types of CBASS systems, types I and II. In fact, the aforementioned technique has shown high sensitivity and specificity rates, except for detection of the cyclase APECO1 gene in the type I CBASS systems. Detection is avoided in this case because 1 out of 13 isolates harbours this gene with several modifications in the region where the crRNA molecule hybridizes, further highlighting the specificity of the LAMP-CRISPR-Cas13a technique.

In summary, the study findings demonstrate the regulatory role of the QS network in the Abi system CBASS. Moreover, we propose an innovative biotechnological application for the LAMP-CRISPR-Cas13a rapid-technique (<2 hours) as an ideal tool for optimizing phage therapy, as it could enable detection of a particular phage-defence mechanism, thus predicting the potential inefficacy of a therapeutic phage. Therefore, the use of the LAMP-CRISPR-Cas13a technique is therefore proposed as a promising means of improving phage treatment selection and patient prognosis.

## AKNOWLEDGEMENTS

We thank the Reference Laboratory, Program for the Prevention and Control of Healthcare-Associated Infections and Antimicrobial Stewardship in Andalucía (PIRASOA, Servicio Andaluz de Salud), for their collaboration.

## AUTHOR CONTRIBUTIONS

C.O-C., and P.F-G., developed the experiments; C.O-C., wrote the original manuscript; L. A., L.B., and D.P-M., collaborated in the developed experiments; I.B., L.F-G, F.F-C and A.B-P analyzed the references as well collaborated in writing manuscript; B.A., and J.C-M., revised the manuscript M.T. supervised all results, validated the work and obtained funding for the research.

